# Chemical exchange saturation transfer for detection of antiretroviral drugs in brain tissue

**DOI:** 10.1101/2021.02.25.432765

**Authors:** Aditya N. Bade, Howard E. Gendelman, JoEllyn McMillan, Yutong Liu

**Affiliations:** Department of Pharmacology and Experimental Neuroscience, University of Nebraska Medical Center, Omaha, NE 68198 USA; Department of Radiology, University of Nebraska Medical Center, Omaha, NE 68198 USA

**Keywords:** CEST, Theranostics, HIV-1, 3TC, Antiretroviral

## Abstract

Human immunodeficiency virus type-1 (HIV-1) antiretroviral drug (ARV) theranostics facilitates biodistribution and efficacy of therapies designed to target viral reservoirs. To this end, we have now deployed intrinsic drug chemical exchange saturation transfer (CEST) contrast to detect ARV distribution within the central nervous system (CNS).

**Methods:** CEST effects for lamivudine (3TC) and emtricitabine (FTC) were measured by asymmetric magnetization transfer ratio analyses in solutions. CEST magnetic resonance imaging (MRI) was performed on 3TC-treated mice with analysis made by Lorentzian fitting.

**Results:** CEST effects of 3TC and FTC hydroxyl and amino protons linearly correlated to drug concentrations. 3TC was successfully detected in brain sub-regions by MRI. The imaging results were validated by measurements of CNS drug concentrations.

**Conclusion:** CEST contrasts can be used to detect ARVs using MRI. Such detection can be used to assess spatial-temporal drug biodistribution. This is most notable within the CNS where drug biodistribution may be more limited with the final goal of better understanding ARV-associated efficacy and potential toxicity.

## Introduction

Although antiretroviral therapy (ART) can prolong the life of human immunodeficiency virus type-1 (HIV-1) infected patients [1, 2], neurocognitive disorders persist and occur in up to 50% of infected patients. Disease ranges from asymptomatic neurocognitive impairment (ANI) to mild neurocognitive disorder (MND) and to the most severe form HIV-associated dementia (HAD) [3]. Persistent viral replication in the central nervous system (CNS) elicits neuroimmune activation and is associated with cognitive decline [4–6]. Thus, it is imperative to begin ART early after infection to slow disease progression and attenuate mental health deficits that are associated with advanced disease [7–11]. ART can be a double edge sword by reversing cognitive decline while in select instances speeding adverse clinical outcomes [12, 13]. The latter include neuropsychiatric, motor and behavioral events [14, 15]. Therefore, the ability to follow drug pharmacokinetics (PK) and biodistribution (BD) could serve as a powerful tool to suppress or perhaps limit the establishment of viral CNS reservoirs and minimize off-target ART effects within the CNS.

Chemical exchange saturation transfer (CEST) as a contrast mechanism was employed previously for drug detection [21]. It arises when an exchangeable proton of a macromolecule is magnetically saturated and transferred to water by chemical exchange causing water MRI signal reduction that reflects macromolecule concentrations. Continuous proton exchanges between the macromolecule and water leading to the buildup of the signal reduction, and the large number of molecules in tissue amplify detection [16–20]. Compared to traditional drug detection systems that tag medicines with an imaging agent or load drugs and imaging agents into a nanoparticle, CEST-based imaging does not require extrinsic drug detection agents. This eliminates the limitations associated with therapeutic efficacy by reduced loading capacity [21] and toxicity [22, 23]. Herein, we developed HIV theranostics based on the CEST contrasts of ARVs. CEST effects of nucleoside reverse transcriptase inhibitors (NRTIs) including lamivudine (3TC) and emtricitabine (FTC) were characterized. Proof-of-concept for CEST was made by injecting 3TC into mice and tracing the drug in brain sub-regions.

## Materials and Methods

### Study approvals

All animal studies were approved by the University of Nebraska Medical Center Institutional Animal Care and Use Committee (IACUC) in accordance with the standards incorporated in the Guide for the Care and Use of Laboratory Animals (National Research Council of the National Academies, 2011).

### Reagents

3TC (lamivudine) was purchased from BOC Sciences (Shirley, NY). FTC (emtricitabine) was purchased from HBCChem (Union City, CA). (Hydroxypropyl)methyl cellulose (HPMC), polysorbate 80 (TWEEN® 80), and lipopolysaccharide (LPS) were purchased from Sigma-Aldrich (St. Louis, MO). Gibco™ DPBS, LC-MS grade water and methanol were purchased from Fisher Scientific (Waltham, MA). 0.9% Sodium Chloride Injection, USP was purchased from Hospira (Lake Forest, IL).

### CEST contrasts of 3TC and FTC

CEST contrasts of 3TC and FTC were measured in saline at 37°C on a 7 Tesla scanner (Bruker PharmaScan 70/16, Billerica, MA) containing a Bruker quadrature RF coil. CEST data was acquired using a Rapid Imaging with Refocused Echoes (RARE) sequence with TR/TE = 4000/42 ms, RARE factor = 16, saturation RF power = 3.6 *μ*T, and duration = 3 s. To construct Z-spectra, saturation frequencies were set from −8 to +8 ppm, step = 0.2 ppm. A second RARE data with saturation RF power = 0.5 *μ*T, and frequencies = −1 to +1 ppm were acquired for B0 correction using WASSR [24]. An image of RF power = 0 *μ*T was acquired as baseline (*S_0_*) for the normalization of CEST images. Following B0 correction and normalization, Z-spectra were constructed. A Z-spectrum is the water signal as a function of saturation frequency, and asymmetric magnetism transfer ratio (*MTR_asym_*) was calculated from the Z-spectrum:

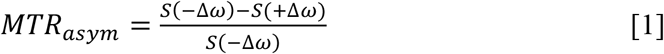

where *Δω* is the frequency offset between the saturation frequency and the water center frequency (0 ppm), and *S*(−*Δω*) and *S*(+*Δω*) are the water signal intensities at the offset frequency lower than water frequency (upfield) and higher than water frequency (downfield), respectively.

### 3TC administration to mice

Male C57BL/6 mice (14-16 weeks old) were purchased from the Jackson Laboratory (Bar Harbor, ME). Mice were randomly distributed in two groups, control and 3TC treated. First, to mimic the inflammatory response of HIV-1 infected patients, mice from both groups (3TC and control) were administered daily 1 mg/kg of lipopolysaccharide (LPS) by intraperitoneal injection in 100 μL sterile saline [25–27] for five days, starting at 24 hours before the first dose of 3TC. The last dose (5^th^ dose) of LPS was administered at 24 hours before MRI imaging at day 5. Second, 3TC solution prepared in a vehicle (0.2% hydroxypropylmethyl cellulose and 0.1% Tween 80 in sterile water) was administered to mice (3TC group) daily by oral gavage for 5 days at a dose of 250 mg/kg. Six hours post-final 3TC dose administration at day 5, mice were scanned for CEST imaging. Control mice were given vehicle alone.

### Animal CEST MRI

CEST imaging was performed on controls (n = 8) and 3TC-treated mice (n = 7) on a 7 Tesla MRI scanner (Bruker BioSpec 70/20, Billerica, MA). A Bruker-made volume quadrature RF coil was employed for signal transmission and a Bruker 4-element coil array was used for signal reception. Respiration and body temperature were monitored during scanning. CEST data was acquired using a RARE sequence (TR/TE = 1600/16 ms, RARE factor = 8) with a continuous RF for saturation with the power = 2 *μ*T, duration = 1 s, saturation frequencies = −5 to 5 ppm in steps of 0.2 ppm. A second CEST data with saturation RF power = 0.5 *μ*T, and frequencies = −1 to +1 ppm were acquired for B0 inhomogeneity correction using WASSR [24]. An image of RF power = 0 *μ*T was acquired as baseline (S_0_) for the normalization of CEST images.

Based on the CEST effects of 3TC shown in Figure 1, the *in vivo* Z-spectra were first normalized using *S*_0_ (baseline image of RF power = 0 *μ*T) and then fitted using a five-pool Lorentzian model of bulk water, aliphatic nuclear Overhauser effect (NOE), magnetization transfer (MT) contrast, amide and amino CEST effects:

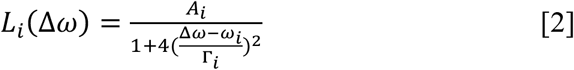

where Δ*ω* is the saturation frequency related to water resonance; *A_i_*, *ω_i_*, and *Γ_i_* are respectively the amplitude, chemical shift and full width at half maximum (FWHM) of the *i*th CEST peak. The initial values of *A_i_*, *ω_i_*, and *Γ_I_* of NOE, amide and MT were set according to previous brain CEST studies [28–34]. The initial values of amino effect were set at *ω* = 2 ppm, *Γ* = 1.5 ppm based on the measurements of 3TC shown in Figure 1. A number of amplitudes (A = 0.05 ~ 0.2) were tried and the fitting results were similar. The integral under each Lorentzian line was calculated pixel by pixel.

**Figure 1.**
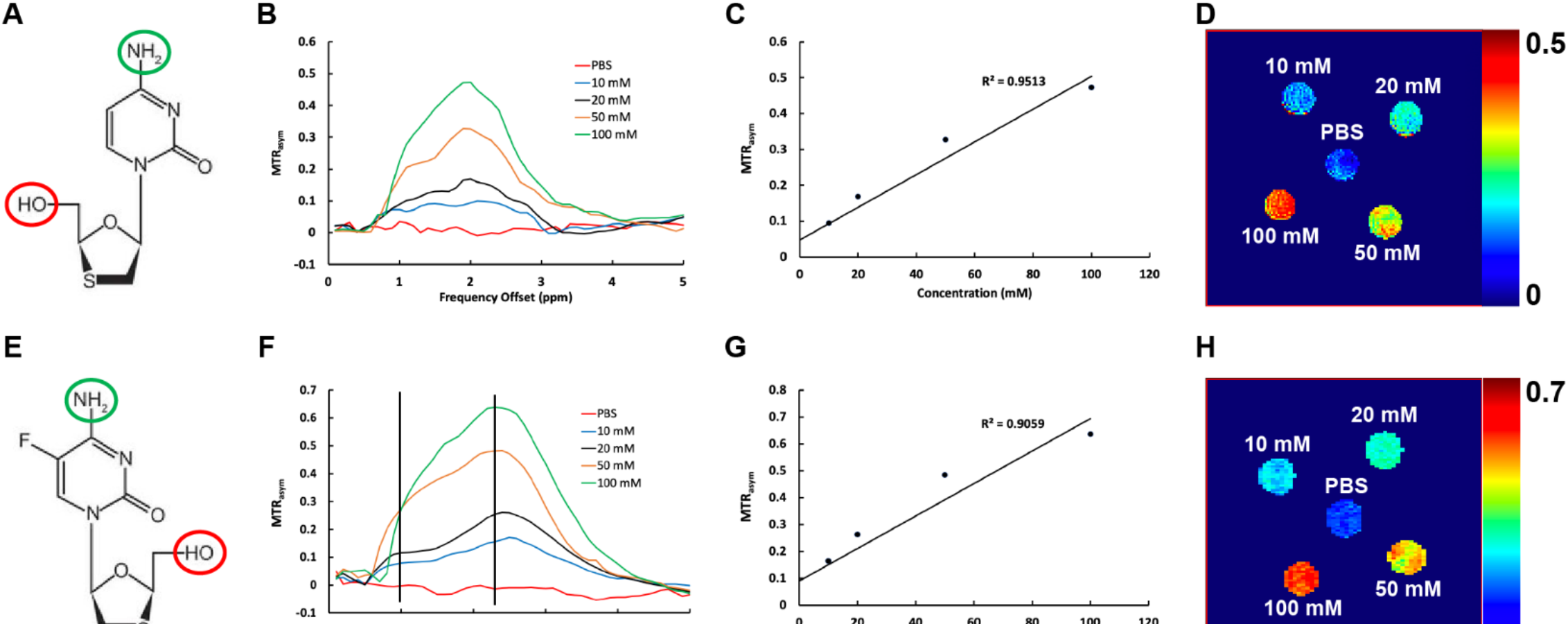
CEST effects of 3TC and FTC. (**A**) Chemical structure of 3TC. The hydroxyl group is enclosed in red circle, and the amino group is enclosed in blue circle. (**B**) *MTR* plots of 3TC and PBS (control) at 37°C. *MTR* increases at 1 ppm and 2 ppm with the 3TC concentration. PBS did not show CEST effect. (**C**) 3TC amino proton CEST effect (MTR@2ppm) increases linearly with 3TC concentration (R^2^ = 0.95). (**D**) Pixel-by-pixel heatmaps of 3TC samples. Color intensity increases with 3TC concentration. The color bar for the heatmaps is at the side of the figure represents *MTR* values. (**E**) Chemical structure of FTC. The hydroxyl group is enclosed in red circle, and the amino group is enclosed in blue circle. (**F**) *MTR* plots of FTC and PBS (control) at 37°C. *MTR* increases at 1 ppm and 2 ppm with the FTC concentration. PBS did not show CEST effect. (**G**) FTC amino proton CEST effect (MTR@2ppm) increases linearly with FTC concentration (R^2^ = 0.90). (**H**) Pixel-by-pixel heatmaps of FTC samples. Color intensity increases with FTC concentration. The color bar for the heatmaps is at the side of the figure represents *MTR* values.

### PK and BD of 3TC in mice

3TC concentrations in plasma and brain tissue samples of 3TC-treated mice (n = 5) were measured by ultraperformance tandem mass spectrometry (UPLC-MS/MS) using a Waters ACQUITY H-class UPLC (Waters, Milford, MA) connected to a Xevo TQ-S micro mass spectrometer [35–37]. All solvents for sample processing and UPLC-MS/MS analysis were LC-MS-grade (Fisher Scientific). 3TC levels in plasma were measured at 6 hours and day 5 time points. Blood samples were collected into heparinized tubes by cheek puncture (submandibular vein) using a 5 mm lancet (MEDIpoint, Inc., Mineola, NY). Collected blood samples were centrifuged at 2,000 × g for 8 minutes to collect plasma. Plasma samples were stored at −80 °C for further quantitation of 3TC levels. At day 5 following MRI imaging, animals were humanely euthanized; and brain regions (hippocampus, cortex, and mid brain) were isolated for quantitation of 3TC concentrations. For plasma drug quantitation 25 μl of plasma was added to 1 ml of ice-cold acetonitrile. Tissue samples were weighed and homogenized in a solution of 90% methanol/10% water. 100 *μ*L of each tissue homogenate was then added to 1 mL of ice-cold methanol. For plasma and tissue drug analysis, acetonitrile-precipitated plasma and methanol-precipitated brain tissue were vortexed for 3 minutes, followed by 10-minute centrifugation at 16000 x g. The resulting supernatant was collected into a new tube and dried down using a SpeedVac (Savant SPD1010, Thermo Scientific). Samples were reconstituted in 80% methanol and 3TC levels were quantitated using UPLC-MS/MS as described [37].

### Statistics

Statistical analyses were conducted using GraphPad Prism 7.0 software (La Jolla, CA). Results from *in vivo* studies were expressed as mean ± SEM. Student’s t tests were performed to compare *in vivo* CEST imaging results from the control and 3TC groups. Pearson’s correlation was used to determine the association between 3TC imaging results and brain tissue drug levels measured by UPLC-MS/MS.

## Results

### CEST contrasts of 3TC and FTC

The CEST contrasts of 3TC and FTC are shown in Figure 1. 3TC is a cytidine analog, and its chemical structure includes a hydroxyl proton and an amino proton (Figure 1A). The CEST contrasts of these were assessed using 3TC solutions at 10, 20, 50, and 100 mM. The *MTR_asym_* plots of 3TC and PBS are shown in Figure 1B. The CEST effects of the hydroxyl and amino protons were observed at 1 and 2 ppm on the *MTR_asym_* plots. The CEST effects were increased with the concentration of 3TC. PBS did not show any CEST effect (Figure 1B). The CEST effect of the amino proton (MTR@2ppm) was linearly proportional to the 3TC concentration with a correlation coefficient R^2^ = 0.95. The amino effects of 3TC samples (MTR@2ppm) are presented in heatmaps in Figure 1D. The color intensity increased with the concentration of 3TC. Concentration-dependent CEST analysis was also performed for the hydroxyl proton using MTR@1pmm. Similarly, the CEST effect of hydroxyl proton was linearly increased with 3TC concentration (R^2^ = 0.94; Supplementary Figure 1A - B). Further, the CEST effects of cytidine analogs were validated using another ARV – FTC (Figure 1E to 1H). Similar to 3TC, concentration dependent CEST effects of the hydroxyl and amino protons of FTC (10, 20, 50 or 100 mM) were observed at 1 and 2.4 ppm, respectively, on the *MTRasym* plot (Figure 1F). However, the CEST effect of FTC amino proton shifted to 2.4 ppm and its magnitude was higher compared to 3TC (Figure 1B vs. Figure 1F). The CEST effect of the amino proton (MTR@2.4ppm) was linearly proportional to the FTC concentration with a correlation coefficient R^2^ = 0.90 (Figure 1G). Increase in color intensity with FTC concentration compared to control (PBS) confirmed the CEST contrast generated by amino group of FTC (Figure 1H). The CEST effect of the hydroxyl (MTR@1ppm) of FTC also linearly increased with the drug concentration (R^2^ = 0.89; Supplementary Figure 1C – D).

### CEST in 3TC-treated mice

CEST effects of 3TC were measured in C57BL/6 male mice brain sub-regions following daily oral drug administration for 5 days at a dose of 250 mg/kg. This dose was about 5 times that of human dosage after animal equivalent dose (AED) calculation [38]. Vehicle to dissolve 3TC was used as control to avoid any confounders. T_2_-weighted images were used as an anatomical reference for brain region-of-interest (ROI) analysis (Figure 2A). The CEST 3TC effect was measured in five sub-regions of the CNS - hippocampus (HIP), cortex (CTX), piriform cortex (PIR), thalamus (TH) and hypothalamus (HY). The Z-spectra of brain regions were built from CEST MRI data. Figure 2B shows representative Z-spectra on HIP of a control and a 3TC-treated mouse. Z-spectra on other brain regions (CTX, TH, PIR, and HY) are shown in Supplementary Figure 2. In a Z-spectrum, the water proton signal is plotted as a function of saturation frequency, and a CEST effect is represented as a signal drop on the Z-spectrum at certain saturation frequency. The evident signal drop from the direct saturation of the bulk water occurred at 0 ppm (Figure 2B). Signal drops at ~ 2 ppm and 3.5 ppm resulted from amino and amide protons, respectively (Figure 2B). The signal at 2 ppm in the 3TC mouse was larger than in the control mouse, indicating the CEST effect of the 3TC amino proton. The signal drop within −1.0 ~ −4.0 ppm resulted from aliphatic NOE (Figure 2B), which is a type of cross-relaxation pathway where spin polarization exchange takes place. A five-pool Lorentzian function was used to fit the bulk water, aliphatic NOE, MT, amino and amide protons. MT contrast is a broad, non-specific signal drop in the Z-spectrum from semi-solid macromolecules [29, 34, 39–42]. Representative fitting results in the control and the 3TC-treated mouse on HIP region are shown in Figure 2C and 2D. The fitting results on CTX, PIR, TH and HY are demonstrated in Supplementary Figure 3. For better visualization, fitted bulk water and MT were removed and fitted functions of NOE, amino and amide protons are shown in Figure 2E and Supplementary Figure 3. Compared to the controls, increase (in both amplitude and linewidth) in amino proton effect (at about 2 ppm) was observed on all brain sub-regions of the 3TC group (Figure 2E and Supplementary Figure 3). The amide proton effects (at about 3.5 ppm) were comparable between the control and the 3TC groups (Figure 2E and Supplementary Figure 3). NOEs were reduced in 3TC-treated mice compared to those of controls. Pixel-by-pixel heatmaps of the integrals of the fitted amino Lorentzian line on each brain sub-region were shown in Figure 3A-3D. T_2_-weighted images were used as an anatomical reference (Figure 3A and 3C). Higher color intensity of the amino CEST effect compared to controls is seen on all brain sub-regions of the 3TC group (Figure 3B and 3D). The 3TC group had a significant increase of amino CEST effect compared to the controls on CTX (p = 0.028), HIP (p = 0.047), TH (p = 0.039), PIR (p = 0.044) and HY (p = 0.024) (Figure 3E). No significant differences were found in amide effects between the control and 3TC groups (Supplementary Figure 4A - E). A trend of decrease was observed for NOE in 3TC mice on HIP (p = 0.099) and TH (p = 0.082) compared to controls (Supplementary Figure 4F - J). This change indicated that 3TC induced biochemical changes in brain.

**Figure 2.**
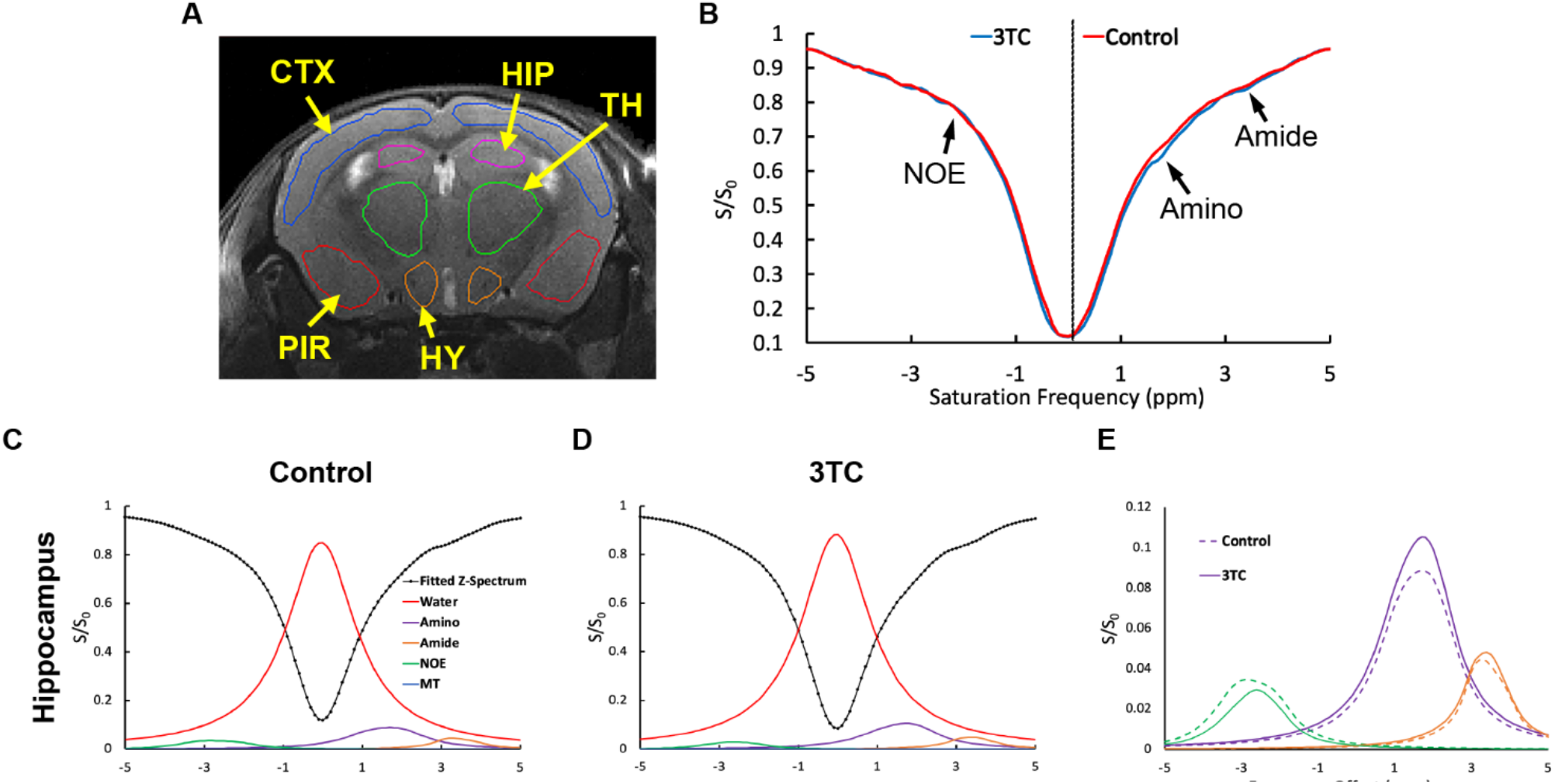
*In vivo* CEST effects of 3TC. (**A**) Regional CEST effects were measured on hippocampus (HIP), cortex (CTX), piriform cortex (PIR), thalamus (TH) and hypothalamus (HY) (**B**) Representative Z-spectra on HIP of a control and a 3TC-treated mouse. (**C**) and (**D**) Representative fitted five Lorentzian functions of bulk water, aliphatic NOE, MT, amino and amide protons on HIP from a control and a 3TC-treated mouse. The sum of the fitted functions is also shown as fitted Z-spectrum. The Lorentzian functions are shown upside down. (E) The fitted NOE, amino and amide protons are shown for better visualization.

**Figure 3.**
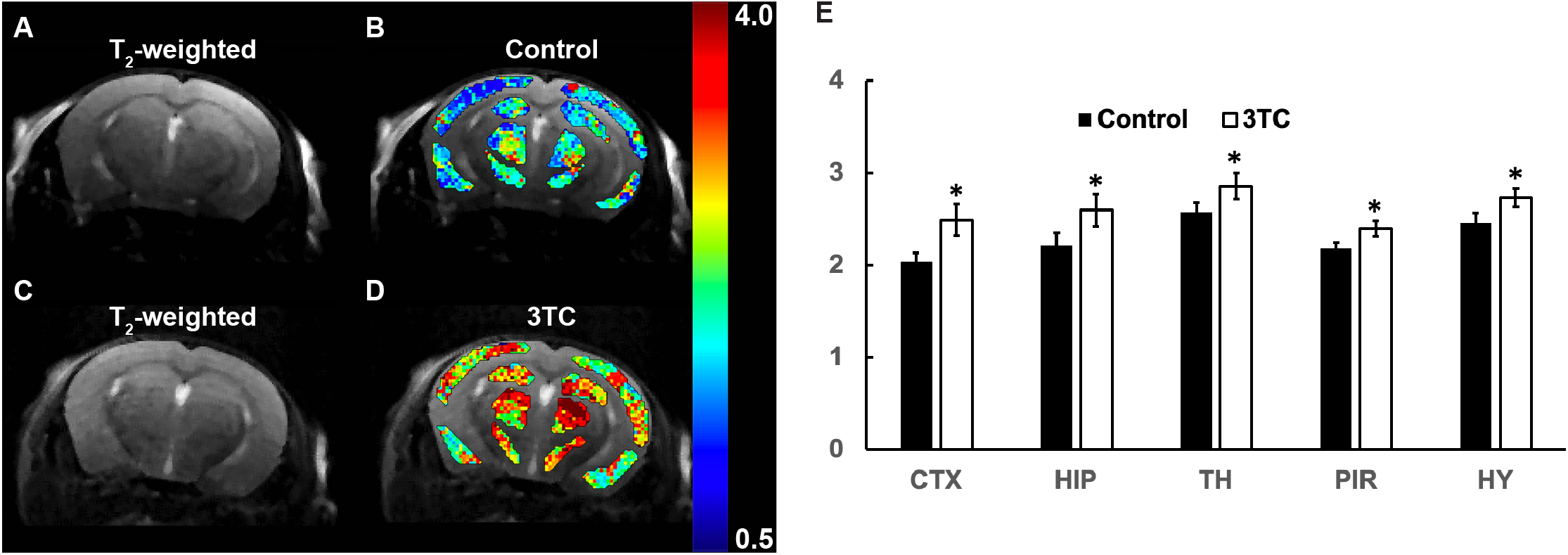
(**A** and **B**) T_2_-weighted image of a control mouse, and pixel-by-pixel Lorentzian line integral heatmaps of fitted amino proton effect superimposed on the respective T_2_-weighted image. (**C** and **D**) T_2_-weighted image of a 3TC-treated mouse, and pixel-by-pixel Lorentzian line integral heatmaps of fitted amino protons superimposed on the respective T_2_-weighted image. (**E**) Group comparisons of fitted integrals of amino proton CEST effect. Student’s t test (two-tailed) was used to compare *in vivo* CEST imaging results from the control and 3TC groups, *: p < 0.05. Data are expressed as mean ± SEM; N = 8 (Control) and N = 7 (3TC)

Further, 3TC levels were measured in plasma and brain sub-regions using UPLC-MS/MS and were correlated to CEST MRI results (Figure 4A and 4B). An average of 1009 ± 154.6 ng/mL and 2819.1 ± 880.1 ng/mL of 3TC was measured in plasma at 6 hours and day 5, respectively. In addition, 3TC levels were 473.3 ± 199.6 ng/g in HIP, 532.6 ± 192 ng/g in CTX and 614.5 ± 227.2 ng/g in mid-brain at day 5. The correlation coefficients of 3TC CEST and UPLC-MS/MS data were R^2^ = 0.62 on CTX, and R^2^ = 0.20 on HIP (Figure 4C and 4D). The correlation of CEST contrasts of CTX and HIP together with 3TC levels in both regions was R^2^ = 0.37 (Figure 4E).

**Figure 4.**
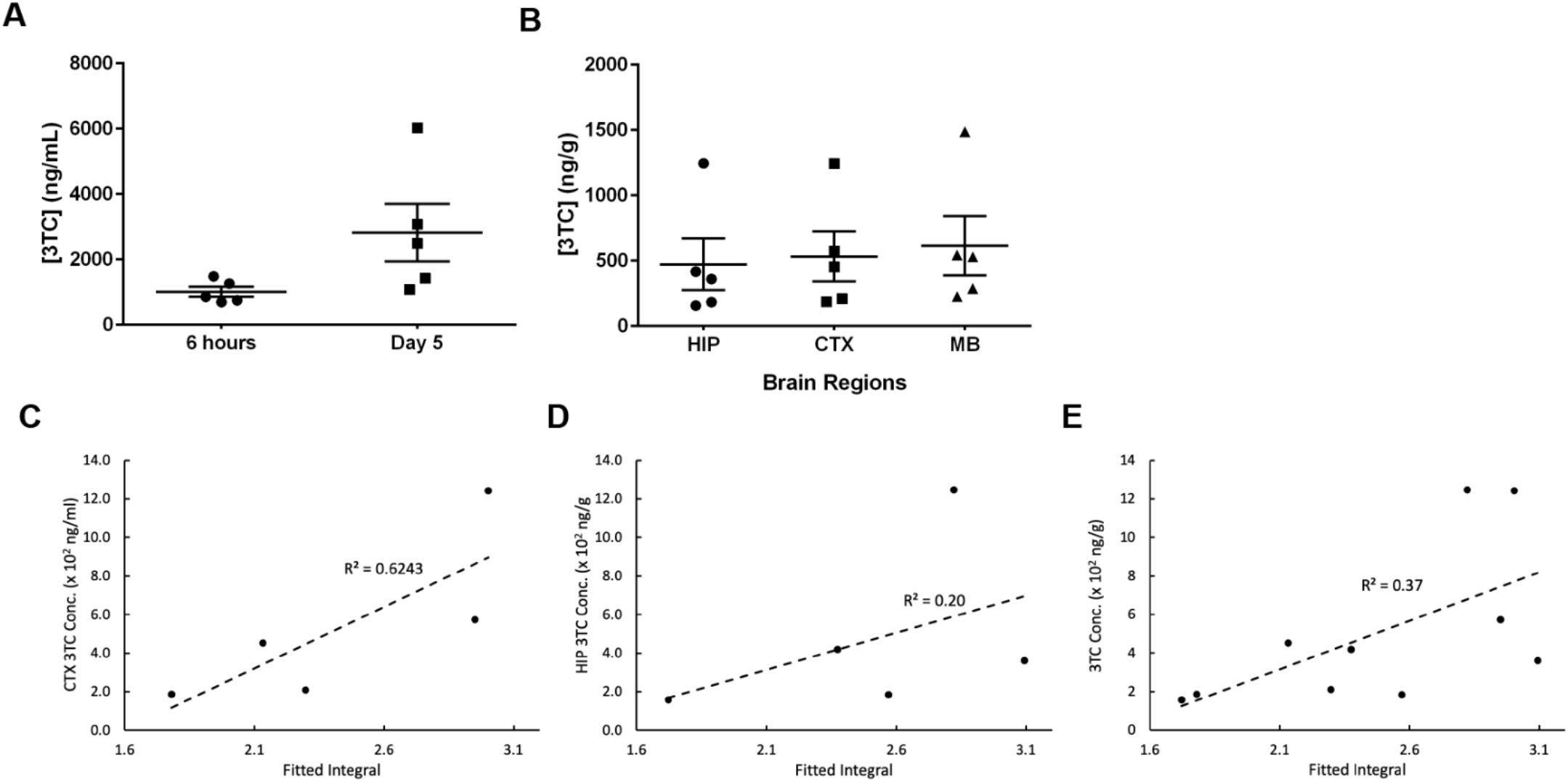
(**A**) Plasma 3TC levels. (B) 3TC concentrations in hippocampus (HIP), cortex (CTX), and mid-brain (MB) at day 5. For both plasma and brain tissue, 3TC concentrations are expressed as mean ± SEM; N = 5/group. (**C**) Correlations of amino proton integrals and tissue 3TC concentrations on CTX. (**D**) Correlations of amino proton integrals and tissue 3TC concentrations on HIP. (**E**) Correlations of amino proton integrals and tissue 3TC concentrations on CTX and HIP combined.

## Discussion

CEST contrasts of ARVs can be characterized and utilized for measurement of biodistribution in brain sub-regions using MRI. In traditional theranostic technologies, drug molecules are tagged with imaging agents or loaded with imaging agents into one nanoparticle. Paramagnetic metals are commonly used for MRI [43–45] and radioactive materials used for PET and SPECT [46–49]. The limitations of the methods are obvious. First, the loading rate of nanoparticles are usually limited to achieve effective therapy and/or imaging sensitivity [21]. Second, toxicity associated with imaging agents and nanoparticles has to be addressed [50–52]. Third, blood brain barrier penetration needs to be considered when designing traditional theranostic methods for the CNS. Unlike the traditional methods, CEST contrasts emerge from the exchangeable protons of drug molecules, and therefore no extrinsic chemical agent is needed for drug imaging. This eliminates the limitations associated with imaging agents used in traditional techniques. Thus, theranostics based on CEST is promising tool for basic and clinical investigations [53].

The development of CEST drug evaluation is possible for drugs that contain slow to intermediate exchanging protons [20, 54, 55]. In the current study, we focused on the assessment of CEST contrasts of 3TC and FTC. Both ARVs are cytidine analogs and possess a hydroxyl proton and an amino proton in the chemical structure. We posit that other NRTIs can, also be detected using parallel CEST MRI method as they also contain hydroxyl and amino protons. For example, tenofovir is an adenosine analog that has an amino group and two hydroxyl groups and abacavir (ABC) is a guanosine analog that has an amino group, an amide group and a hydroxyl group. Notably, CEST also has the potential to image other classes of ARVs such as integrase strand transfer inhibitors (INSTIs), non-nucleoside reverse transcriptase inhibitors (NNRTIs) and protease inhibitors provided that have slow to intermediate exchanging protons. For ARVs from these classes, characterization of CEST effects can be completed by structural evaluation, and exchange rate calculations [56, 57].

Although we tested the CEST-based brain imaging, the method with a few modifications in the Lorentzian fitting algorithm can be utilized on other tissues such as liver, kidney and spleen. This significantly expands the applications of the CEST-based detection techniques. Moreover, direct *in vivo* imaging of ARVs could enable long term PK and BD studies, which is critical for the development of long-acting injectables of ARVs. It is now well accepted that long-acting ARVs could affect drug adherence and as such reduce viral transmission, prevent new infections, and limit the emergence of viral drug resistance [58–61]. There is an increasing need for personalized ARV regimens to provide efficient treatment while minimizing toxicity and decreasing the risk of viral resistance developing [62–64]. HIV theranostics that measures real-time tissue drug levels will help the design of personalized treatments tailored for individual patients.

A successful bioimaging technique measures the biodistribution of an ARV with sensitivity and specificity. In this study, the relatively high correlations between imaging results and UPLC-MS/MS measurements of 3TC were observed. This showed that CEST imaging is sensitive to quantify drug levels in CNS. However, the 3TC dose used in this study was about five times that of the clinical dose. Moreover, LPS used to induce inflammation as occurs in HIV-1 patients could enhance brain penetration of 3TC. These technical limitations will be addressed by testing the MRI method on various dosages of ARVs in HIV-1 infected humanized mice. Such approach will help to determine and improve the sensitivity of the CEST contrasts. The saturation RF power and duration in the current study was selected based on previous *in vivo* CEST studies [40, 65]. RF parameters will be further optimized in the future using ARVs in cells as suggested in [41] to improve sensitivity for *in vivo* environments.

The major challenge for sufficient specificity for *in vivo* ARV measurements is the contaminations by protons from background biomolecules including proteins and metabolites. Additionally, the MT contrasts from semisolid macromolecules and NOE of macromolecules also confound the ARV CEST contrasts [66]. Simple asymmetric *MTR* analysis, usually, fails for *in vivo* CEST data analysis. Thus, more advanced data analysis methods like the Lorentzian line-shape fitting needs be used to improve the analysis outcome [17, 41, 67–70]. This study demonstrated that the Lorentzian function successfully detected 3TC *in vivo* by fitting the amino proton CEST effect. We posit that the specificity of 3TC detection can be further improved by fitting the combined CEST effects of the amino and hydroxyl protons. Based on this idea, we are extending the Lorentzian method to a multiple-peak algorithm that fits CEST effects of different protons simultaneously in an ARV using Lorentzian functions and polynomials. This method is inspired by previous work [28, 40, 71], and can be further expanded to identify different ARVs used in combined ARV regimens.

In summary, we demonstrated that the CEST effects of cytidine analogs (3TC and FTC) can be characterized. These characterized properties can be harnessed for *in vivo* drug detection and quantification using advanced data analysis algorithms like Lorentzian line-shape fitting. ARV theranostics based on intrinsic CEST effects will advance the field in evaluating the spatial-temporal biodistribution of drugs and in understanding ARV-associated efficacy and toxicity.

## Abbreviations

3TC: lamivudine
ABC: abacavir
AED: animal equivalent dose
ANI: asymptomatic neurocognitive impairment
ARV: antiretroviral
ART: antiretroviral therapy
BBB: blood-brain barrier
BD: biodistribution
CEST: chemical exchange saturation transfer
CNS: central nervous system
CTX: cortex
FTC: emtricitabine
FWHM: full width at half maximum
HAD: HIV-associated dementia
HAND: HIV-1-associated neurocognitive disorder
HIP: hippocampus
HIV-1: human immunodeficiency virus type-1
HY: hypothalamus
INSTIs: integrase strand transfer inhibitors
LPS: lipopolysaccharide
MND: mild neurocognitive disorder
MRI: magnetic resonance imaging
MT: magnetization transfer
MTRasym: asymmetric magnetism transfer ratio
NNRTIs: non-nucleoside reverse transcriptase inhibitors
NOE: nuclear Overhauser effect
NPAEs: ARVs-associated neuropsychiatric adverse events
NRTIs: nucleoside reverse transcriptase inhibitors
PET: positron emission tomography
PIR: piriform cortex
PK: pharmacokinetics
RARE: Rapid Imaging with Refocused Echoes
RF: radiofrequency
ROI: region-of-interest
SPECT: single photon emission computed tomography
TH: thalamus
TVF: tenofovir

## Acknowledgements

Authors thank Bhagya Laxmi Dyavar Shetty for technical assistance with drug quantitation by UPLC-MS/MS, and Melissa Mellon and Lirong Xu in MRI core facility for mouse imaging. This study was partially supported by Nebraska Research Initiative.

## References

1. Phanuphak, N. and R.M. Gulick, HIV treatment and prevention 2019: current standards of care. Curr Opin HIV AIDS, 2020. 15(1): p. 4–12.

2. Ghosn, J., et al., HIV. The Lancet, 2018. 392(10148): p. 685–697.

3. Valcour, V., et al., Central nervous system viral invasion and inflammation during acute HIV infection. J Infect Dis, 2012. 206(2): p. 275–82.

4. Saylor, D., et al., HIV-associated neurocognitive disorder--pathogenesis and prospects for treatment. Nature reviews. Neurology, 2016. 12(4): p. 234–248.

5. Hong, S. and W.A. Banks, Role of the immune system in HIV-associated neuroinflammation and neurocognitive implications. Brain Behav Immun, 2015. 45: p. 1–12.

6. Katuri, A., et al., Role of the inflammasomes in HIV-associated neuroinflammation and neurocognitive disorders. Experimental and Molecular Pathology, 2019. 108: p. 64–72.

7. Van den Hof, M., et al., CNS penetration of ART in HIV-infected children. J Antimicrob Chemother, 2018. 73(2): p. 484–489.

8. Carvalhal, A., et al., Central nervous system penetration effectiveness of antiretroviral drugs and neuropsychological impairment in the Ontario HIV Treatment Network Cohort Study. J Neurovirol, 2016. 22(3): p. 349–57.

9. Decloedt, E.H., et al., Central nervous system penetration of antiretroviral drugs: pharmacokinetic, pharmacodynamic and pharmacogenomic considerations. Clin Pharmacokinet, 2015. 54(6): p. 581–98.

10. Yilmaz, A., R.W. Price, and M. Gisslen, Antiretroviral drug treatment of CNS HIV-1 infection. J Antimicrob Chemother, 2012. 67(2): p. 299–311.

11. Nwogu, J.N., et al., Pharmacokinetic, Pharmacogenetic, and Other Factors Influencing CNS Penetration of Antiretrovirals. AIDS Res Treat, 2016. 2016: p. 2587094.

12. Asiedu, N., I. Kretchy, and E. Asampong, Psycho-behavioral factors associated with neurocognitive performance among people living with HIV on antiretroviral therapy in Accra, Ghana. Afr Health Sci, 2020. 20(2): p. 487–596.

13. Mellgren, Å., et al., Longitudinal trends and determinants of patient-reported side effects on ART-a Swedish national registry study. PLoS One, 2020. 15(12): p. e0242710.

14. Treisman, G.J. and O. Soudry, Neuropsychiatric Effects of HIV Antiviral Medications. Drug Saf, 2016. 39(10): p. 945–57.

15. Abers, M.S., W.X. Shandera, and J.S. Kass, Neurological and psychiatric adverse effects of antiretroviral drugs. CNS Drugs, 2014. 28(2): p. 131–45.

16. Dou, W., et al., Chemical exchange saturation transfer magnetic resonance imaging and its main and potential applications in pre-clinical and clinical studies. Quant Imaging Med Surg, 2019. 9(10): p. 1747–1766.

17. McMahon, M.T. and A.A. Gilad, Cellular and Molecular Imaging Using Chemical Exchange Saturation Transfer. Top Magn Reson Imaging, 2016. 25(5): p. 197–204.

18. Pankowska, A., et al., Chemical exchange saturation transfer (CEST) as a new method of signal obtainment in magnetic resonance molecular imaging in clinical and research practice. Pol J Radiol, 2019. 84: p. e147–e152.

19. Sinharay, S. and M.D. Pagel, Advances in Magnetic Resonance Imaging Contrast Agents for Biomarker Detection. Annu Rev Anal Chem (Palo Alto Calif), 2016. 9(1): p. 95–115.

20. Wu, B., et al., An overview of CEST MRI for non-MR physicists. EJNMMI Phys, 2016. 3(1): p. 19.

21. Dreifuss, T., et al., A challenge for theranostics: is the optimal particle for therapy also optimal for diagnostics? Nanoscale, 2015. 7(37): p. 15175–84.

22. Rahman, M., et al., Role of Graphene Nano-Composites in Cancer Therapy: Theranostic Applications, Metabolic Fate and Toxicity Issues. Curr Drug Metab, 2015. 16(5): p. 397–409.

23. Desai, N., Challenges in development of nanoparticle-based therapeutics. Aaps j, 2012. 14(2): p. 282–95.

24. Kim, M., et al., Water saturation shift referencing (WASSR) for chemical exchange saturation transfer (CEST) experiments. Magn Reson Med, 2009. 61(6): p. 1441–50.

25. Banks, W.A., et al., Lipopolysaccharide-induced blood-brain barrier disruption: roles of cyclooxygenase, oxidative stress, neuroinflammation, and elements of the neurovascular unit. Journal of Neuroinflammation, 2015. 12(1): p. 223.

26. Maggioli, E., et al., Estrogen protects the blood-brain barrier from inflammation-induced disruption and increased lymphocyte trafficking. Brain Behav Immun, 2016. 51: p. 212–222.

27. Haruwaka, K., et al., Dual microglia effects on blood brain barrier permeability induced by systemic inflammation. Nature Communications, 2019. 10(1): p. 5816.

28. Chen, L., et al., Investigation of the contribution of total creatine to the CEST Z-spectrum of brain using a knockout mouse model. NMR Biomed, 2017. 30(12).

29. Debnath, A., et al., Glutamate-Weighted CEST Contrast After Removal of Magnetization Transfer Effect in Human Brain and Rat Brain with Tumor. Mol Imaging Biol, 2020.

30. Goerke, S., et al., Relaxation-compensated APT and rNOE CEST-MRI of human brain tumors at 3 T. Magn Reson Med, 2019. 82(2): p. 622–632.

31. Singh, A., et al., Evaluating the feasibility of creatine-weighted CEST MRI in human brain at 7 T using a Z-spectral fitting approach. NMR Biomed, 2019. 32(12): p. e4176.

32. Windschuh, J., et al., Correction of B1-inhomogeneities for relaxation-compensated CEST imaging at 7 T. NMR Biomed, 2015. 28(5): p. 529–37.

33. Zaiss, M., et al., Chemical exchange saturation transfer MRI contrast in the human brain at 9.4T. Neuroimage, 2018. 179: p. 144–155.

34. Cai, K., et al., CEST signal at 2ppm (CEST@2ppm) from Z-spectral fitting correlates with creatine distribution in brain tumor. NMR Biomed, 2015. 28(1): p. 1–8.

35. Guo, D., et al., Creation of a Long-Acting Nanoformulated 2’,3’-Dideoxy-3’-Thiacytidine. J Acquir Immune Defic Syndr, 2017. 74(3): p. e75–e83.

36. Smith, N., et al., A long acting nanoformulated lamivudine ProTide. Biomaterials, 2019. 223: p. 119476.

37. Soni, D., et al., Synthesis of a long acting nanoformulated emtricitabine ProTide. Biomaterials, 2019. 222: p. 119441.

38. Nair, A.B. and S. Jacob, A simple practice guide for dose conversion between animals and human. Journal of basic and clinical pharmacy, 2016. 7(2): p. 27–31.

39. Bagga, P., et al., In vivo GluCEST MRI: Reproducibility, background contribution and source of glutamate changes in the MPTP model of Parkinson’s disease. Scientific Reports, 2018. 8(1): p. 2883.

40. Chen, L., et al., Creatine and phosphocreatine mapping of mouse skeletal muscle by a polynomial and Lorentzian line-shape fitting CEST method. Magn Reson Med, 2019. 81(1): p. 69–78.

41. Zaiss, M., et al., Inverse Z-spectrum analysis for spillover-, MT-, and T1 -corrected steady-state pulsed CEST-MRI--application to pH-weighted MRI of acute stroke. NMR Biomed, 2014. 27(3): p. 240–52.

42. Zhuang, Z., et al., Mapping the Changes of Glutamate Using Glutamate Chemical Exchange Saturation Transfer (GluCEST) Technique in a Traumatic Brain Injury Model: A Longitudinal Pilot Study. ACS Chemical Neuroscience, 2019. 10(1): p. 649–657.

43. Dadfar, S.M., et al., Iron oxide nanoparticles: Diagnostic, therapeutic and theranostic applications. Adv Drug Deliv Rev, 2019. 138: p. 302–325.

44. Lux, F., et al., Gadolinium-based nanoparticles for theranostic MRI-radiosensitization. Nanomedicine (Lond), 2015. 10(11): p. 1801–15.

45. Zhu, L., et al., Magnetic nanoparticles for precision oncology: theranostic magnetic iron oxide nanoparticles for image-guided and targeted cancer therapy. Nanomedicine (Lond), 2017. 12(1): p. 73–87.

46. Bailly, C., et al., Immuno-PET for Clinical Theranostic Approaches. Int J Mol Sci, 2016. 18(1).

47. Bodet-Milin, C., et al., Clinical Results in Medullary Thyroid Carcinoma Suggest High Potential of Pretargeted Immuno-PET for Tumor Imaging and Theranostic Approaches. Front Med (Lausanne), 2019. 6: p. 124.

48. Lenzo, N.P., D. Meyrick, and J.H. Turner, Review of Gallium-68 PSMA PET/CT Imaging in the Management of Prostate Cancer. Diagnostics (Basel), 2018. 8(1).

49. Pruis, I.J., G. van Dongen, and S.E.M. Veldhuijzen van Zanten, The Added Value of Diagnostic and Theranostic PET Imaging for the Treatment of CNS Tumors. Int J Mol Sci, 2020. 21(3).

50. Gupta, N., et al., A Review of Theranostics Applications and Toxicities of Carbon Nanomaterials. Curr Drug Metab, 2019. 20(6): p. 506–532.

51. Bruna Galdorfini, C.-A., et al., Drug Delivery Using Theranostics: An Overview of its Use, Advantages and Safety Assessment. Current Nanoscience, 2020. 16(1): p. 3–14.

52. Li, M., et al., Preclinical evaluation of <sup>203/212</sup>Pb-based theranostics-dosimetry and renal toxicity.<strong/>. Journal of Nuclear Medicine, 2020. 61(supplement 1): p. 289–289.

53. Li, Y., et al., CEST theranostics: label-free MR imaging of anticancer drugs. Oncotarget, 2016. 7(6): p. 6369–78.

54. Sherry, A.D. and Y. Wu, The importance of water exchange rates in the design of responsive agents for MRI. Curr Opin Chem Biol, 2013. 17(2): p. 167–74.

55. Soesbe, T.C., Y. Wu, and A. Dean Sherry, Advantages of paramagnetic chemical exchange saturation transfer (CEST) complexes having slow to intermediate water exchange properties as responsive MRI agents. NMR Biomed, 2013. 26(7): p. 829–38.

56. Dixon, W.T., et al., A concentration-independent method to measure exchange rates in PARACEST agents. Magn Reson Med, 2010. 63(3): p. 625–32.

57. Sun, P.Z., et al., Quantitative chemical exchange saturation transfer (qCEST) MRI--RF spillover effect-corrected omega plot for simultaneous determination of labile proton fraction ratio and exchange rate. Contrast media & molecular imaging, 2014. 9(4): p. 268–275.

58. Dash, P.K., et al., Long-acting nanoformulated antiretroviral therapy elicits potent antiretroviral and neuroprotective responses in HIV-1-infected humanized mice. Aids, 2012. 26(17): p. 2135–44.

59. Dash, P.K., et al., Sequential LASER ART and CRISPR Treatments Eliminate HIV-1 in a Subset of Infected Humanized Mice. Nat Commun, 2019. 10(1): p. 2753.

60. Canetti, D. and V. Spagnuolo, An evaluation of cabotegravir for HIV treatment and prevention. Expert Opin Pharmacother, 2020: p. 1–12.

61. Flexner, C., et al., LONG-ACTING DRUGS AND FORMULATIONS FOR THE TREATMENT AND PREVENTION OF HIV. Int J Antimicrob Agents, 2020: p. 106220.

62. Mu, Y., et al., The dawn of precision medicine in HIV: state of the art of pharmacotherapy. Expert opinion on pharmacotherapy, 2018. 19(14): p. 1581–1595.

63. Cusato, J., et al., Precision medicine for HIV: where are we? Pharmacogenomics, 2018. 19(2): p. 145–165.

64. Lengauer, T., N. Pfeifer, and R. Kaiser, Personalized HIV therapy to control drug resistance. Drug Discovery Today: Technologies, 2014. 11: p. 57–64.

65. Chen, L., et al., High-resolution creatine mapping of mouse brain at 11.7 T using non-steady-state chemical exchange saturation transfer. NMR in Biomedicine, 2019. 32(11): p. e4168.

66. Pagel, M.M., The Pursuit of Theranostics with CEST MRI. Theranostics, 2016. 6(10): p. 1601–1602.

67. Kim, J., et al., A review of optimization and quantification techniques for chemical exchange saturation transfer MRI toward sensitive in vivo imaging. Contrast media & molecular imaging, 2015. 10(3): p. 163–178.

68. Liu, G., et al., Nuts and bolts of chemical exchange saturation transfer MRI. NMR in biomedicine, 2013. 26(7): p. 810–828.

69. Liu, Y., et al., Novel HIV Detection by Chemical Exchange Saturation Transfer (CEST), in 2019 World Molecular Imaging Congress. 2019, World Molecular Imaging Society: Montreal, Canada.

70. Vinogradov, E., A.D. Sherry, and R.E. Lenkinski, CEST: from basic principles to applications, challenges and opportunities. J Magn Reson, 2013. 229: p. 155–72.

71. Ryoo, D., et al., Detection and Quantification of Hydrogen Peroxide in Aqueous Solutions Using Chemical Exchange Saturation Transfer. Anal Chem, 2017. 89(14): p. 7758–7764.

