## Supplemental Figures for "Chemical exchange saturation transfer for detection of antiretroviral drugs in brain tissue"

### Supplementary Information

#### Supplementary Figures

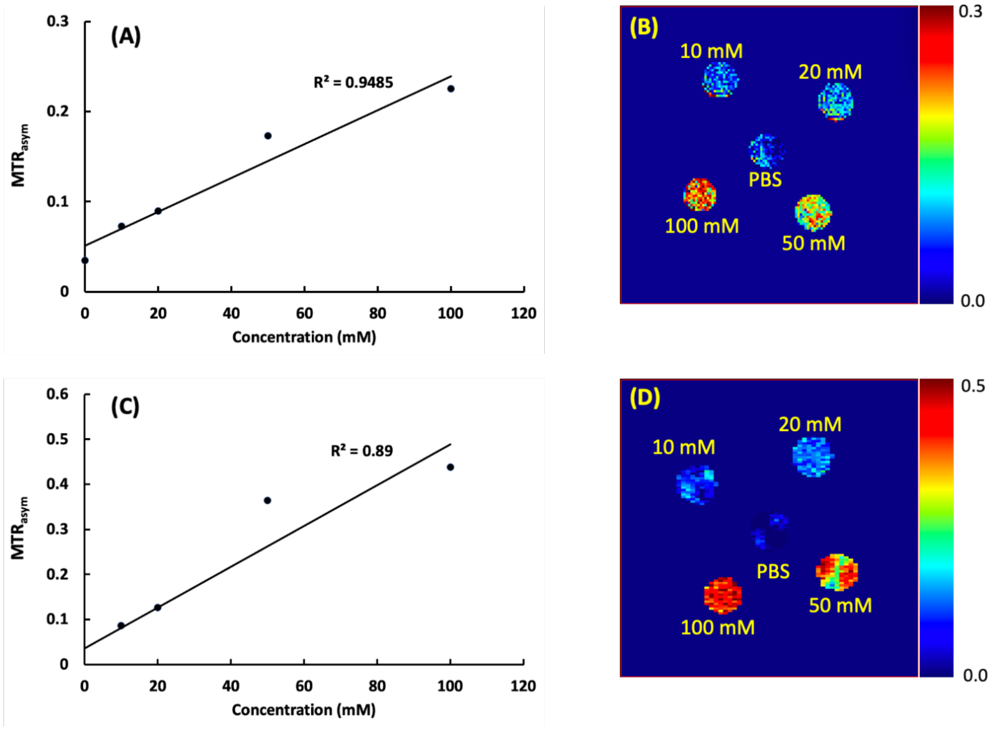


**Supplementary Figure 1.** Hydroxyl proton CEST effects of 3TC and FTC. (**A**) 3TC hydroxyl proton CEST effect (MTR@1ppm) increases linearly with 3TC concentration ($R^2$ = 0.95). (**B**) Pixel-by-pixel heatmaps of 3TC samples. Color intensity increases with 3TC concentration. The color bar for the heatmaps is at the side of the figure represents MTR values. (**C**) FTC hydroxyl proton CEST effect (MTR@1ppm) increases linearly with FTC concentration ($R^2$ = 0.89). (**D**) Pixel-by-pixel heatmaps of FTC samples. Color intensity increases with FTC concentration. The color bar for the heatmaps is at the side of the figure represents MTR values.

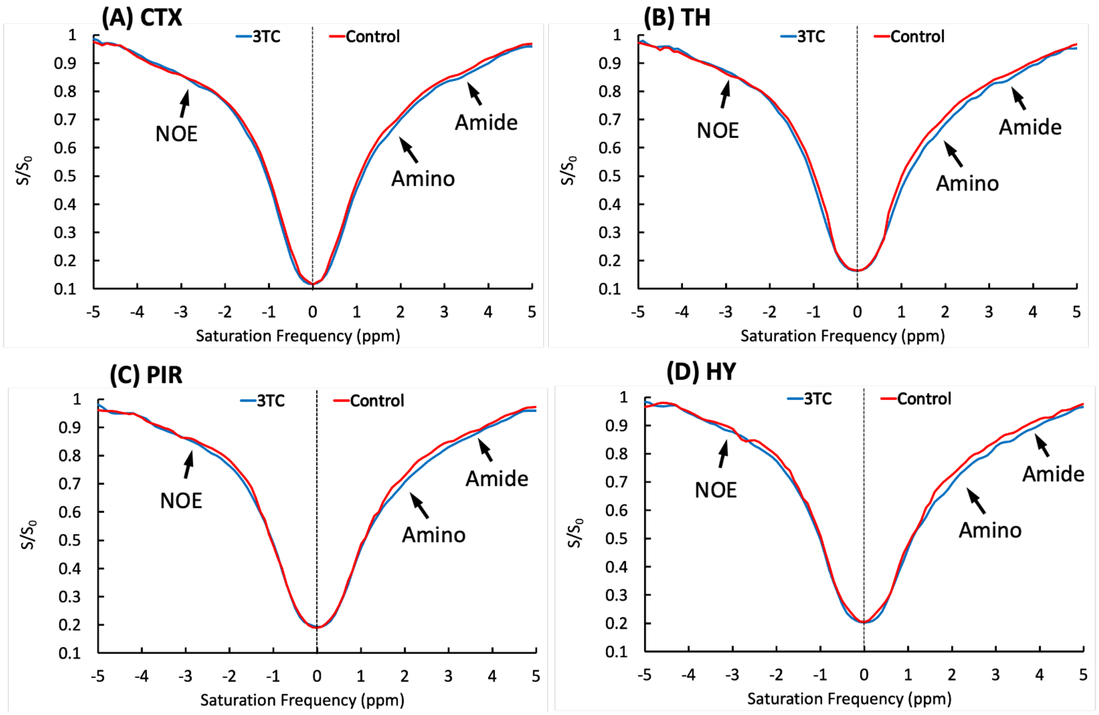


**Supplementary Figure 2.** Representative Z-spectra of a control and a 3TC-treated mouse on (**A**) CTX (cortex), (**B**) TH (thalamus), (**C**) PIR (piriform cortex), and (D) HY (hypothalamus). The effects of NOE, amnio and amide protons are indicated by arrows.

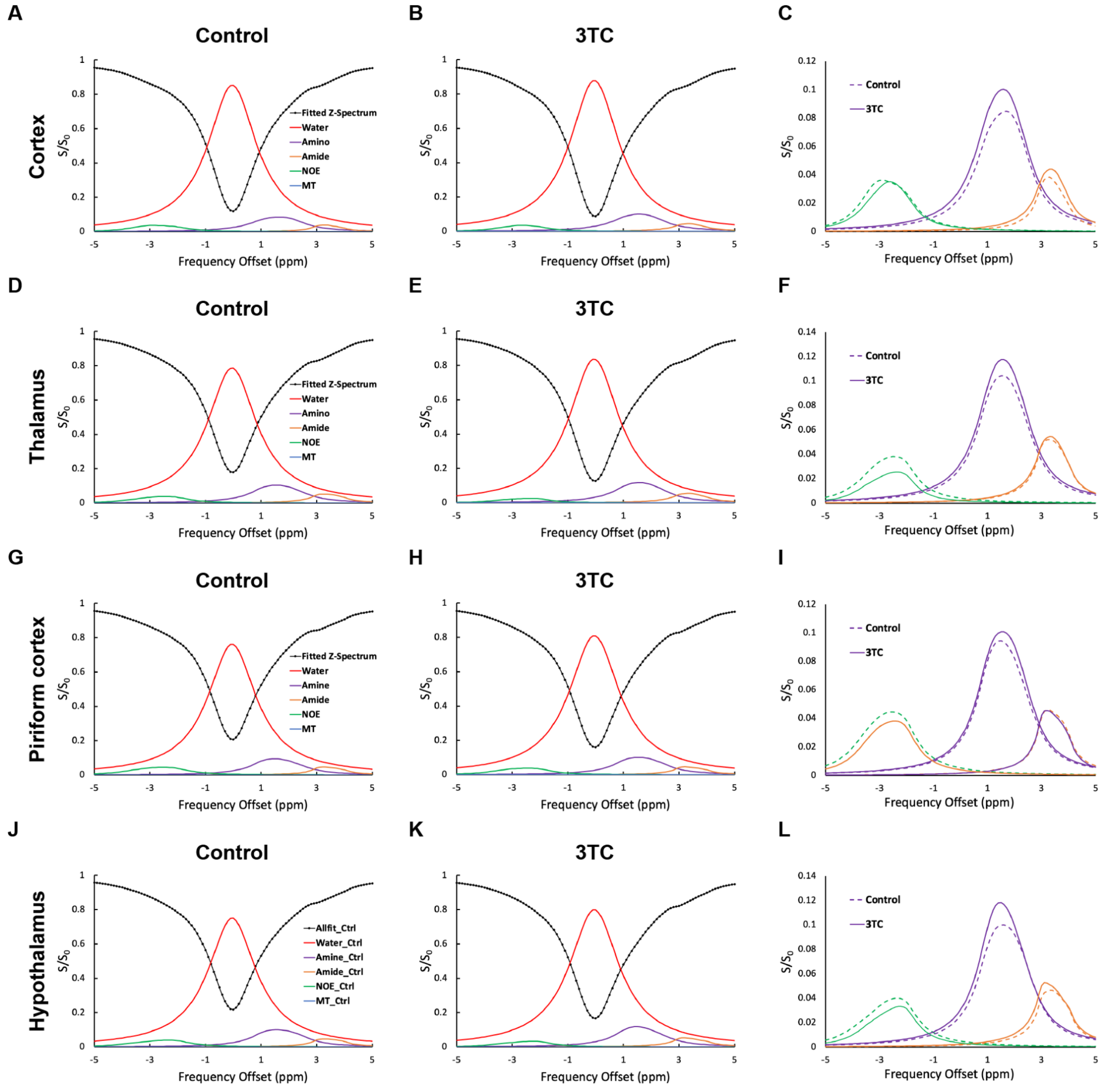


**Supplementary Figure 3.** Representative fitted five Lorentzian functions of bulk water, aliphatic NOE, MT, amino and amide protons from a control and a 3TC-treated mouse on (**A and B**) CTX (cortex), (**D and E**) TH (thalamus), (**G and H**) PIR (piriform cortex), and (**J and K**) HY (hypothalamus). The sum of the fitted functions is shown as fitted Z-spectrum. The Lorentzian functions are shown upside down. The fitted NOE, amino and amide proton CEST effects on (**C**) CTX, (**F**) TH, (**I**) PIR and (**L**) HY are shown for better visualization.

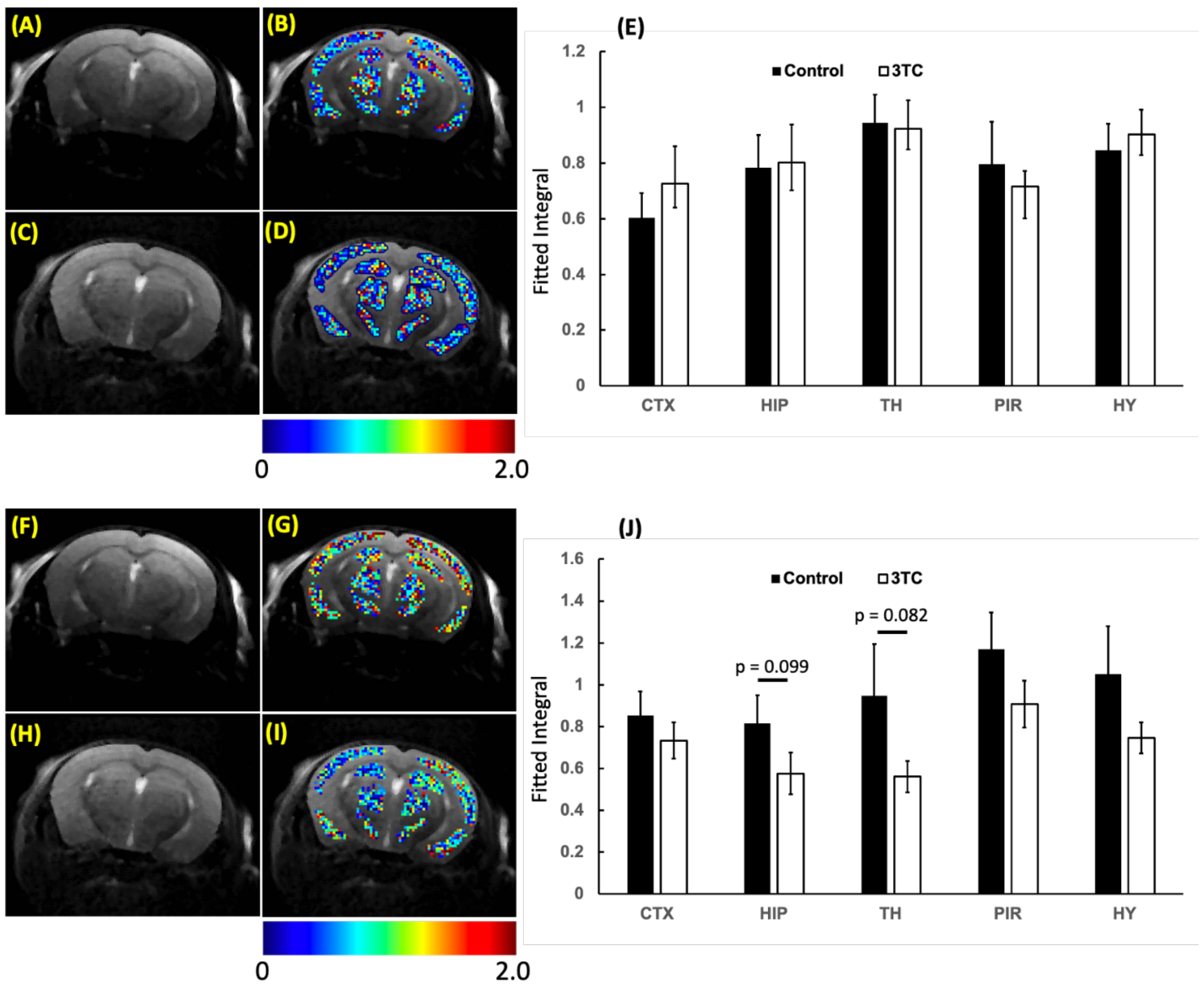


**Supplementary Figure 4.** (**A** and **B**) $T_2$-weighted image of a control mouse, and pixel-by-pixel Lorentzian line integral heatmaps of fitted amide proton effect superimposed on the respective $T_2$-weighted image. (**C** and **D**) $T_2$-weighted image of a 3TC-treated mouse, and pixel-by-pixel integral maps of fitted amide effect superimposed on the respective $T_2$-weighted image. (**E**) Group comparisons of fitted integrals of amide proton CEST effect. (**F** and **G**) $T_2$-weighted image of a control mouse, and pixel-by-pixel Lorentzian line integral heatmaps of fitted NOE superimposed on the respective $T_2$-weighted image. (**H** and **I**) $T_2$-weighted image of a 3TC-treated mouse, and pixel-by-pixel integral maps of NOE superimposed on the respective $T_2$-weighted image. (**J**) Group comparisons of fitted integrals of NOE. For both amide proton and NOE comparisons, Student's t test (two-tailed) was used to compare *in vivo* CEST imaging results from the control and 3TC groups. Data are expressed as mean ± SEM; N = 8 (Control) and N = 7 (3TC)
